# Intracranial Mapping of Response Latencies and Task Effects for Spoken Syllable Processing in the Human Brain

**DOI:** 10.1101/2024.04.05.588349

**Authors:** Vibha Viswanathan, Kyle M. Rupp, Jasmine L. Hect, Emily E. Harford, Lori L. Holt, Taylor J. Abel

## Abstract

Prior lesion, noninvasive-imaging, and intracranial-electroencephalography (iEEG) studies have documented hierarchical, parallel, and distributed characteristics of human speech processing. Yet, there have not been direct, intracranial observations of the latency with which regions *outside the temporal lobe* respond to speech, or how these responses are impacted by task demands. We leveraged human intracranial recordings via stereo-EEG to measure responses from diverse forebrain sites during (i) passive listening to /bi/ and /pi/ syllables, and (ii) active listening requiring /bi/-versus-/pi/ categorization. We find that neural response latency increases from a few tens of ms in Heschl’s gyrus (HG) to several tens of ms in superior temporal gyrus (STG), superior temporal sulcus (STS), and early parietal areas, and hundreds of ms in later parietal areas, insula, frontal cortex, hippocampus, and amygdala. These data also suggest parallel flow of speech information dorsally and ventrally, from HG to parietal areas and from HG to STG and STS, respectively. Latency data also reveal areas in parietal cortex, frontal cortex, hippocampus, and amygdala that are not responsive to the stimuli during passive listening but are responsive during categorization. Furthermore, multiple regions—spanning auditory, parietal, frontal, and insular cortices, and hippocampus and amygdala—show greater neural response amplitudes during active versus passive listening (a task-related effect). Overall, these results are consistent with hierarchical processing of speech at a macro level and parallel streams of information flow in temporal and parietal regions. These data also reveal regions where the speech code is stimulus-faithful and those that encode task-relevant representations.

**New & Noteworthy:** We leverage direct, intracranial electroencephalography recordings to document speech information flow through diverse sites in the human forebrain, including areas where reported electrode coverage has historically been limited. Our data are consistent with hierarchical processing of speech at a macro level and parallel streams of information flow in temporal and parietal regions. They also reveal regions in the auditory pathway where stimulus-faithful speech codes are transformed to behaviorally relevant representations of speech content.

## Introduction

Anatomical tract tracing studies suggest that several different pathways may be available for auditory and speech processing (Dick and Tremblay, 2012). A robust literature exists discussing hierarchical, parallel, and distributed characteristics of auditory and speech information flow through human cortex (McClelland et al., 2006; Ahveninen et al., 2006; Hickok and Poeppel, 2007; Rauschecker and Scott, 2009; Bizley and Cohen, 2013; Kemmerer, 2014; Hamilton et al., 2021). Invasive work in non-human primates and noninvasive magnetoencephalography (MEG) and electroencephalography (EEG) studies in humans suggest that primary (core) auditory cortex is the first cortical site for sound processing, while secondary and other auditory cortical areas (belt and parabelt) have longer initial-response latencies than the core (Kaas et al., 1999). However, intracranial recordings in humans suggest that there may also be parallel information processing pathways, with a superior temporal gyrus (STG) zone activating in parallel with some sites in primary auditory cortex (Heschl’s gyrus; HG) (Nourski et al., 2014; Hamilton et al., 2021). Prior intracranial electrophysiological studies on auditory processing have focused mainly on early auditory cortical areas. Indeed, direct, intracranial measurements of speech information flow through forebrain regions outside of early auditory areas are limited.

Besides understanding the trajectories of neural information flow, another foundational question for models of speech processing is how stimulus-faithful speech codes in the ascending auditory pathway are transformed to behaviorally relevant representations of speech content. Task-related auditory processing has been studied in diverse cortical sites in both human and animal models (Hall et al., 2000; Fritz et al., 2007; Nourski et al., 2017). Consistent with the view that primary auditory cortex represents earlier stages of sound processing, studies using non-human primates suggest that core auditory areas have a relatively simple receptive-field organization, while belt and parabelt areas are increasingly more complex in their response properties (Rauschecker and Scott, 2009). In line with this, non-primary but not primary auditory cortex in humans shows sound encoding that is invariant to background noise (Kell and McDermott, 2019), and has specializations for voice (Belin et al., 2000) and abstract speech units (e.g., at the phonemic, semantic, and lexical levels) versus simply encoding speech acoustics (Davis and Johnsrude, 2003; Okada et al., 2010; Keshishian et al., 2023). However, both animal studies (Fritz et al., 2003; Atiani et al., 2009; Niwa et al., 2012) and noninvasive studies in humans [using functional magnetic resonance imaging (fMRI; Jäncke et al., 1999; De Martino et al., 2015) and MEG (Woldorff et al., 1993; Kauramäki et al., 2012; Ding and Simon, 2012)] suggest that task-dependent modulation of neural responses may be present as early as primary auditory cortex (HG).

Historically, our understanding of speech information flow was derived from lesion studies in patients suffering acute brain injury (e.g., from stroke or head trauma; Wernicke, 1874). More recently, studies have leveraged noninvasive imaging techniques such as fMRI, EEG, and MEG. Although these techniques benefit from whole-brain coverage, they are generally limited by their relatively poor joint spatio-temporal resolution (Vazquez and Noll, 1998; Samuelsson et al., 2021). Invasive recordings in animals provide high spatio-temporal specificity but are constrained by the extent to which they can inform models of human speech processing (Petkov et al., 2009). Human intracranial EEG (iEEG) in patients with epilepsy provides better spatial specificity than noninvasive scalp EEG. However, traditional iEEG recordings with electrocorticography (ECoG) grid electrodes cover only superficial (i.e., on the pial surface of the cortex) gyral regions like STG but do not include deeper regions such as HG and STS (Mercier et al., 2022). More recently, some human intracranial studies used precisely implanted depth electrodes to examine speech information flow and task-related processing in deeper structures, including HG (Nourski et al., 2014; Nourski et al., 2017; Hamilton et al., 2021; Keshishian et al., 2023). However, prior human intracranial reports of speech latency have been restricted to the auditory cortex, with no reported electrode coverage outside the temporal lobe. Indeed, the latency with which different brain regions outside the temporal lobe respond to speech sounds or how these responses are impacted by task demands has not been well studied with direct, intracranial measurements.

In the current study, we leveraged intracranial stereo-EEG (sEEG) electrodes implanted in epilepsy surgery patients to document patterns of speech information flow through the human forebrain and how speech-evoked neural responses are influenced by task demands. We directly examined responses to speech syllables across diverse brain regions—including both deep and superficial structures—using sEEG electrodes implanted in HG, STG, STS, insula, inferior frontal gyrus (IFG), and other frontal, parietal, and basal-forebrain sites. We measured neural responses across two different task conditions: (i) passive listening to /bi/ and /pi/ speech syllables, and (ii) active listening that required a /b/ versus /p/ category-identity decision on the same syllables. Across these diverse sites, we quantified neural response latency for the speech syllables and the extent to which response amplitudes differ between passive and active conditions (i.e., task-related effects).

## Materials and Methods

### Participants

Simultaneous sEEG and behavioral data were acquired from six neurosurgical patients with drug-resistant epilepsy as part of clinical evaluation for epilepsy surgery at Children’s Hospital of Pittsburgh. Patients undergoing sEEG electrode implantation to the supratemporal plane who require chronic (i.e., >3 days) sEEG implantation and recording of the temporal lobe for localization of epileptic foci or clinical language mapping were candidates for inclusion. Exclusion criteria included significant medical or neuropsychological impairment, which would have rendered the patients unable to participate in the proposed research. sEEG electrode implantation was planned based on clinical necessity by epileptologists not involved in the research, protecting against research-related conflicts of interest. Participants completed the task in one session during post-surgical monitoring in the hospital. All procedures were approved by the University of Pittsburgh Institutional Review Board, and all participants provided written informed consent or assent (in the case of adolescent 14-to-17-year-olds) to participate prior to sEEG implantation.

Patient demographics information is provided in Table 1. All participants were native speakers of North American English. Where available, language laterality (dominant hemisphere for language) was determined for participants based on documentation in the electronic medical record (EMR) from clinical neurology visits or fMRI lateralization studies. No patient had any record of hearing loss noted in their clinical chart. Moreover, all stimuli were presented at a supra-threshold level that was judged to be loud but comfortable by each subject.

**Table 1.**
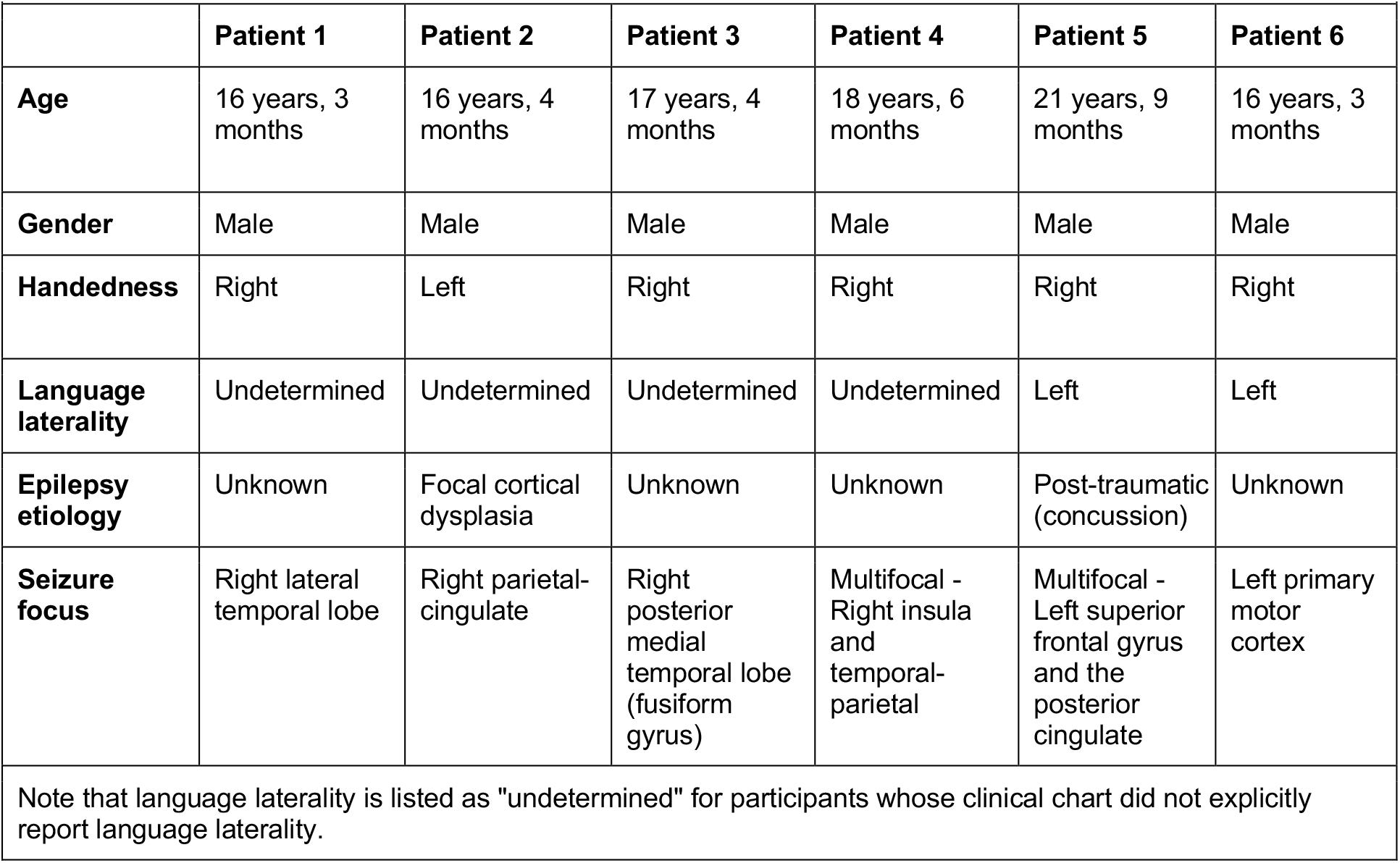
Patient demographics information.

|  | **Patient 1** | **Patient 2** | **Patient 3** | **Patient 4** | **Patient 5** | **Patient 6** |
| --- | --- | --- | --- | --- | --- | --- |
| **Age** | 16 years, 3 months | 16 years, 4 months | 17 years, 4 months | 18 years, 6 months | 21 years, 9 months | 16 years, 3 months |
| **Gender** | Male | Male | Male | Male | Male | Male |
| **Handedness** | Right | Left | Right | Right | Right | Right |
| **Language laterality** | Undetermined | Undetermined | Undetermined | Undetermined | Left | Left |
| **Epilepsy etiology** | Unknown | Focal cortical dysplasia | Unknown | Unknown | Post-traumatic (concussion) | Unknown |
| **Seizure focus** | Right lateral temporal lobe | Right parietal-cingulate | Right posterior medial temporal lobe (fusiform gyrus) | Multifocal - Right insula and temporal-parietal | Multifocal - Left superior frontal gyrus and the posterior cingulate | Left primary motor cortex |

Note that language laterality is listed as "undetermined" for participants whose clinical chart did not explicitly report language laterality.

### sEEG electrode implantation

sEEG electrode implantation was performed as described in earlier work (Abel et al., 2018). Briefly, Dixi Medical Microdeep electrodes were used, with a diameter of 0.8 mm, contact length of 2 mm, and center-to-center spacing of 3.5 mm. Each electrode contained between 8 and 18 contacts. Each such contact will henceforth be referred to as an sEEG channel.

### Behavioral paradigm

Figure 1 illustrates the behavioral paradigm, which was adapted from Hodson et al. (2023). Stimuli consisted of natural utterances of the monosyllabic English words *bee* and *pea* spoken by a female monolingual English speaker that were manipulated to vary across voice onset time (VOT; -10, 0, 10, 20, 30, 40, or 50 ms) and fundamental frequency (F0; 170, 180, 190, 240, 250, or 260 Hz), as in Idemaru and Holt (2011). Each stimulus was normalized to the same root-mean-square (RMS) amplitude.

**Figure 1.**
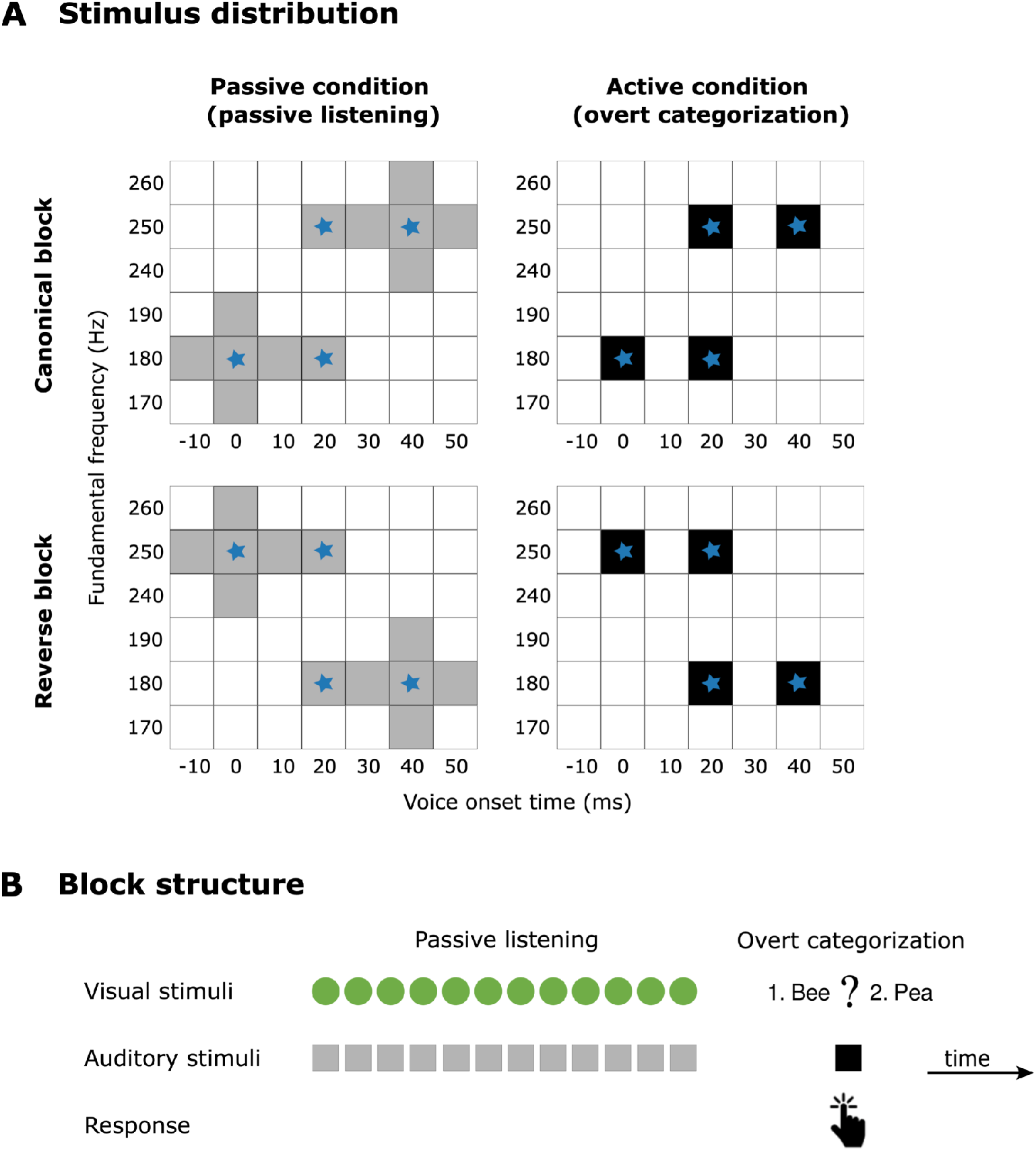
Behavioral paradigm. Panel A shows the range of voice onset time and fundamental frequency values for the *bee*-*pea* monosyllabic-word stimuli used in the study, separately for the passive (gray shaded boxes) and active (black shaded boxes) listening conditions. The stimulus distribution varied according to the experimental block. During passive listening, Canonical blocks exposed subjects to a typical pattern of VOT and F0 covariation in English (shorter VOT and lower F0 values for /b/ and longer VOT and higher F0 values for /p/) and Reverse blocks exposed subjects to a subtle ‘accent’ that reversed this correlation. The stimuli in the active condition consisted primarily of *bee-pea* utterances with a perceptually ambiguous VOT (20 ms); a smaller proportion (40% of total) of stimuli had VOT values that could be unambiguously mapped to either /b/ or /p/ (VOT=0 and 40 ms). The latency analysis used all available stimuli in each condition (active, passive). Blue stars indicate stimuli that were used in the analysis of task-related effects. Panel B shows the structure of each experimental block. Each block consisted of 13 stimuli, of which the first 12 involved passive listening to *bee-pea* speech utterances (Panel A, passive condition); the 13th stimulus required subjects to overtly categorize a stimulus (Panel A, active condition) as either *bee* or *pea* via a button press. Visual cues guided the timing of audio stimulus presentation. The entire experiment had 100 blocks (i.e., 1300 stimuli) in total; the first 50 blocks were Canonical blocks and the rest Reverse blocks.

Note that the experimental paradigm was designed to study neural correlates of adaptive plasticity in speech perception, specifically, how passive exposure to *bee-pea* speech syllables with different acoustic regularities influences overt categorization of *bee-pea* utterances with a perceptually ambiguous VOT (20 ms; see Hodson et al., 2023). In this paradigm, the speech-input regularity either matches a typical pattern of VOT and F0 covariation in English (Canonical blocks; smaller VOT and F0 values for /b/ and larger VOT and F0 values for /p/) or introduces an ‘accent’ (Reverse blocks). Although the current study uses this paradigm—which was designed for a different purpose—here, we focus on speech information flow and examine neural response latency and task effects (contrasting passive and active listening) for syllable processing.

The experiment had 100 blocks, of which the first 50 were Canonical blocks and the rest Reverse blocks. Each block presented 13 250-ms speech stimuli, of which the first 12 involved passive listening (passive condition) to *bee* and *pea* utterances whose VOT and F0 varied across stimuli (as in Figure 1A) and the 13th required overt categorization (active condition) as *bee* or *pea* via a button press. The stimuli in the active condition consisted primarily of *bee-pea* utterances with a perceptually ambiguous VOT (20 ms); 40% of the stimuli in the active condition had VOT values that could be unambiguously mapped to either /b/ or /p/ (VOT=0 and 40 ms).

The inter-stimulus interval (i.e., the time between the end of a given stimulus and the start of the next stimulus) between passive stimuli was 300 ms and that between the final passive stimulus and the active stimulus in each block was 600 ms. Moreover, there was a 300 ms pause at the end of each block after the subjects’ button-press response. The length of the full experimental session was ∼20 min.

Visual cues were provided throughout the experiment (shown in Figure 1). During passive listening, a progress bar indicated the number of stimuli elapsed within the block. The visual stimulus for the 13th (overt categorization) stimulus was a prompt with the words *bee* and *pea* and a question mark. Subjects reported their category-identity decision by tapping a button to indicate which word they heard.

### Stimulus calibration

We used perceptual calibration based on subjects’ loudness rating to ensure clear audibility of all stimuli. Subjects were presented a ∼2-min-long calibration stimulus—which was created by concatenating all the *bee/pea* stimuli—that matched the actual experimental stimuli in both spectrum and RMS. The sound level was adjusted until the subject rated the volume as “loud but comfortable”. The level setting was not adjusted thereafter.

### Hardware and software

A laptop computer controlled all aspects of the experiment, including sound delivery, stimulus onset marking via DAQ pulses, and storing stimulus and experiment information. Audio stimuli were presented diotically through ER-3C (Etymotic, Elk Grove Village, IL) insert earphones coupled to foam ear tips. All audio signals were digitized at a sampling rate of 48 kHz. Neural data were recorded and stored using a separate computer at a sampling rate of 1 kHz using the Nomad (Ripple Neuro, Salt Lake City, UT) 512-channel Neural Interface Processor (model R02000), Grapevine (Ripple Neuro) digital and analogue I/O, and Micro2 front end amplifiers (Ripple Neuro). The audio was fed into a distribution amplifier (Rolls Co., Murray, UT; model DA134); the split audio signal was presented to patients and simultaneously recorded synchronously at a sampling rate of 30 kHz with the sEEG data by the Ripple amplifier. The synchronously recorded audio and DAQ pulses together were used to synchronize the sEEG data with the audio stimuli. Subjects responded with a button box (Response Time Box, v6).

Stimulus presentation was controlled using custom MATLAB (The MathWorks, Inc., Natick, MA) routines. Offline data analyses were performed using custom software in MATLAB. Statistical analyses were performed using R (R Core Team; www.R-project.org).

### Electrode localization

Electrode localization was performed to map the different sEEG channels for each patient to different regions in the Julich brain atlas (Amunts et al., 2020). Cortical surfaces were reconstructed from a preoperative MRI using Freesurfer (Fischl et al., 2004). Using the software package Brainstorm (Tadel et al. 2011), the MRI was then co-registered with a postoperative CT scan, and channels were localized. MNI normalization was performed using Brainstorm’s implementation of SPM12’s nonlinear warping. This MNI deformation field was then used to warp the Julich volumetric atlas into patient space (Amunts et al., 2020; Eickhoff et al., 2005; Evans et al., 2012), and each channel was localized to a brain region by finding the region label of the closest voxel.

### sEEG pre-processing

In line with pre-processing procedures used in prior literature (Mesgarani and Chang, 2012; Mesgarani et al., 2014; Hamilton et al., 2020), the raw sEEG data were re-referenced to the average signal across channels. Then, the data were filtered in the high-gamma band (75-150 Hz) using a linear phase FIR filter (filter order was 68 ms, yielding a transition bandwidth of 15 Hz) and forward-reverse filtering to correct for the filter’s group delay. Because a key goal of the analysis was to estimate response latency in different brain regions, this filtering step to extract high-gamma band neural responses was performed with zero group delay (i.e., non-causal filtering). Note that we restricted our analyses to the high-gamma band because prior literature suggests that high-gamma-band responses reflect local neuronal activity whereas lower frequency neural responses (e.g., in the delta-theta band) reflect synchronized activity across cortical regions and brain networks (Canolty and Knight, 2010; Buzsáki et al., 2012).

The envelope of the sEEG in the high-gamma band (hereafter referred to as high-gamma activity; HGA) was extracted by half-wave rectifying the filter output from the previous step and low-pass filtering the result between 0 and 30 Hz using a linear-phase (i.e., constant group delay) non-negative (to prevent negative values in the output) causal FIR filter that was constructed using a Hanning window of length 33 ms (window length was derived as the ratio of the sEEG sampling rate to the desired filter cutoff). Causal FIR filtering was used for envelope extraction because a zero-group-delay non-causal filter would smear envelopes back in time, leading to systematic under-estimation of onset latency. Because our procedure to estimate latency (see section: Neural response latency analysis) defines latency as the time of initial deviation of the neural response amplitude from the pre-stimulus baseline, the non-zero group delay of the FIR filter we used to extract envelopes would not bias our latency estimates.

The HGA was epoched into stimulus-specific responses using an epoch window starting at -100 ms and ending at 450 ms relative to the audio stimulus onset. The logarithm (base 10) of the result was taken to ensure normally distributed response data (Thomson and Chave, 1991). Then, the neural response in each epoch was baseline corrected by subtracting the mean neural activity in the baseline period (−100 to 0 ms relative to the audio stimulus onset) and dividing the result by the standard deviation (STD) in the baseline period to yield a z-score. Finally, sEEG epochs with average (over time and channels) squared HGA deviating more than four STD from the mean across epochs were discarded.

We considered channels in the temporal, parietal, and frontal lobes, as well as in insula, cingulate, and basal forebrain (e.g., hippocampus, amygdala), as available, in our analyses. We analyzed only those regions for which we had recordings from at least two channels and only those channels that were responsive to the speech stimuli. To find stimulus-responsive channels, we computed the mean evoked response for each channel and condition (passive, active) by averaging the HGA z-score over all epochs and dividing the result by the square root of the number of epochs to normalize the result to z-score units. Only those channels whose post-stimulus-onset neural responses had a mean (over time) evoked response z-score exceeding 1.96 [corresponding to the 95% confidence interval (CI)] in at least four time-resolution windows (the number of time-resolution windows in each epoch is the effective number of time samples after accounting for reduction in time resolution from filtering operations during pre-processing) in either the passive or active condition were retained and used in all further analyses in this study; the remaining channels were discarded. The number of time-resolution windows in each epoch was calculated as the epoch duration post stimulus onset (0.45 s) divided by the reciprocal of the smallest filter cutoff used in analysis (30 Hz); thus, the number of time-resolution windows is ∼13 and the number of measured time samples in each such window is ∼34.

### Neural response latency analysis

We computed neural response latency following audio stimulus onset for each sEEG channel, subject, and task condition as the time point of initial deviation of the pre-processed sEEG HGA from baseline. To compute latency, we averaged the pre-processed HGA for a specific channel, subject, and task condition over all epochs. Then, we computed the mean and 95% confidence interval (CI) for the baseline period (−100 to 0 ms relative to audio stimulus onset). Next, we calculated the earliest time point (T1) at which the mean HGA exceeded the baseline 95% CI (i.e., exceeded baseline mean + 1.96 times standard error of the baseline mean; note that 1.96 corresponds to 95% CI for the distribution under the null hypothesis that there is no stimulus-evoked response) and remained above it for at least 30 ms. Then, we fit a straight line (L1) with data points -5 to 10 ms around T1. Finally, we took the time at which L1 crossed the baseline mean to be the latency. Figure 2 illustrates our latency computation procedure. The combination of our sEEG pre-processing approach and latency computation procedure renders our latency estimates invariant to any group delays and response smearing in time introduced by the filtering process. We report latency for a brain region only if there were at least two responsive channels in that region across subjects.

**Figure 2.**
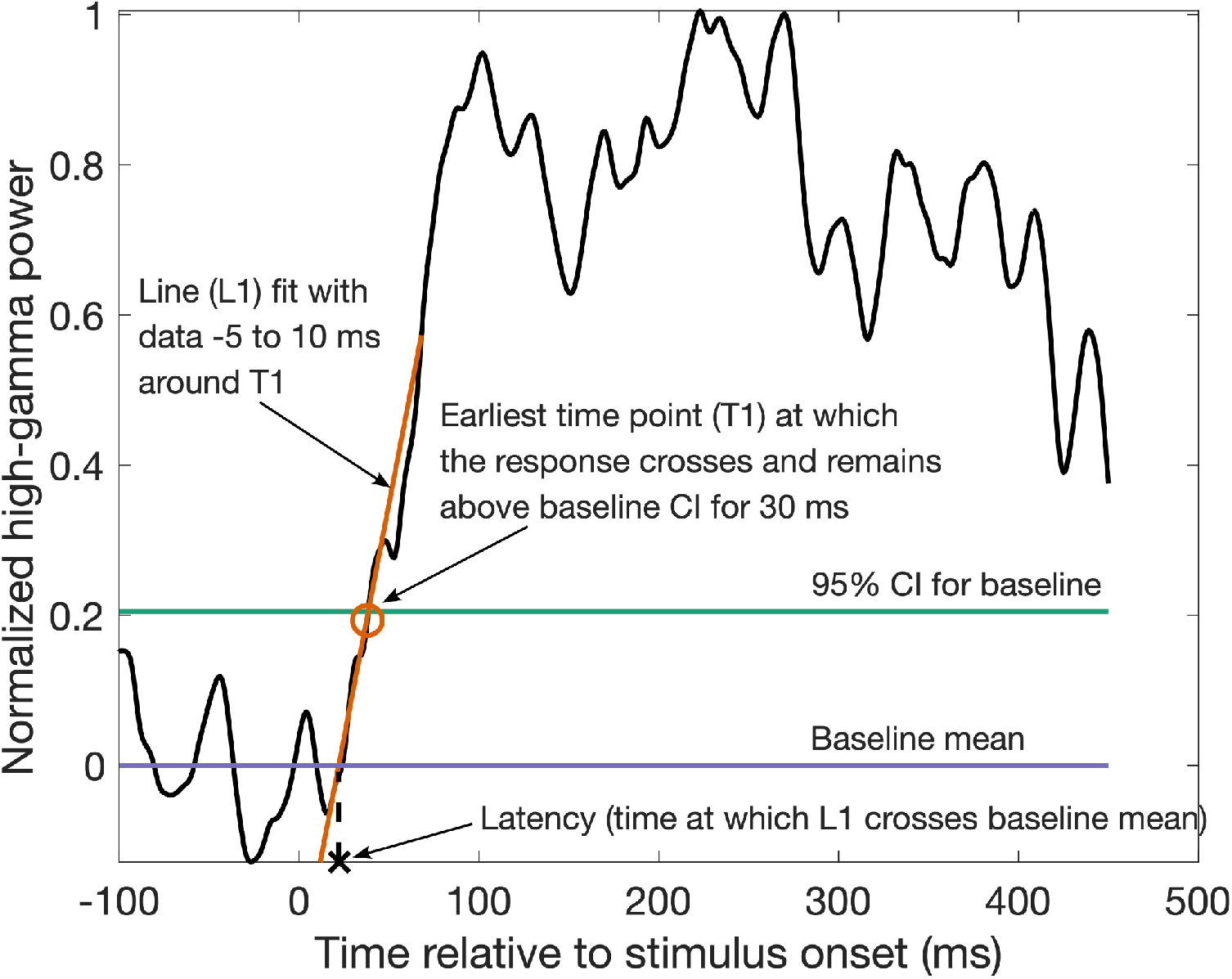
Schematic of latency computation. Latency was computed for each sEEG channel, subject, and task condition (passive, active) from channel-specific HGA as follows. The pre-processed HGA was first averaged over epochs, then the earliest time point (T1; orange circle) following audio stimulus onset at which the mean HGA crossed the baseline 95% confidence interval (CI; green line) and remained above the CI for at least 30 ms was computed. A straight line (L1; orange line) was fit with the set of data points -5 ms before to 10 ms after T1. Finally the time at which L1 crossed the baseline mean (purple line) was taken to be the latency (black cross) of that channel.

### Analysis of task-related effects

To quantify task-related effects for each brain region, we averaged the pre-processed HGA in each condition (passive, active) over just those epochs corresponding to stimuli that were exactly matched between the two conditions and over the time period starting from the channel- and condition-specific latency (if latency was not calculable in a given condition, the latency in the other condition was used instead) and ending at the end of the epoch. We then computed the mean of this result over all region-specific channels across subjects and took the difference in the mean neural response amplitude between the active and passive conditions as the task-related neural response boost.

### Statistical analysis

To test for significant differences in neural response latency across different brain regions, we used a linear mixed-effects model. We fit separate models for right- and left-hemisphere data. Latency in the passive condition in different channels across subjects was the response variable, and brain region (factor variable with seven levels: HG, STG, STS, parietal, frontal, insula, and basal-forebrain) and subject were predictors. Subject effects were treated as random. ANOVA (Type II Wald F tests with Kenward-Roger degree of freedom; Kenward and Roger, 1997) was used for statistical testing.

To test whether the neural response in each brain region shows a significant task-related effect, we used nonparametric permutation testing (Nichols and Holmes, 2002). Under the null hypothesis of zero task effect, the change in neural response amplitude from the passive to the active condition (i.e., the task-related neural response boost) is zero on average. Thus, under the null hypothesis, the passive and active conditions are equivalent and the labels “passive” and “active” can be swapped randomly to generate examples of neural response differences that would be observed under the null hypothesis. This approach was used to generate a separate null distribution for each brain region, pooling data over region-specific sEEG channels. To generate a single realization from each null distribution, the procedure described in the Section “Analysis of task-related effects” was used but after randomly assigning each epoch either the “passive” or “active” label within subject and channel in such a way that there were exactly the same number of epochs in each condition as in the correctly labeled data. This procedure was repeated with 1000 distinct randomizations to generate the full null distribution for each brain region. Finally, for each region, the task-related effect computed for the correctly labeled data was compared with the corresponding null distribution to assign an uncorrected p-value. The Benjamini and Hochberg false-discovery rate (FDR) procedure was used to correct for multiple comparisons across the different brain regions at a 5% FDR (Benjamini and Hochberg, 1995).

### Data availability

The datasets used in the current study are available from T.J.A. on reasonable request.

## Results

### Mapping speech information flow using intracranial neural response latencies

Figure 3 shows single-channel latency of sEEG high-gamma-band envelope responses to speech in the passive condition on a standard MNI brain (for analysis details, see Materials and Methods: Neural response latency analysis). Figure 4 shows mean latency for each brain region (Julich atlas; Amunts et al., 2020) in the passive condition; for those regions for which latency was not calculable in the passive condition (i.e., the latency criteria were not met, indicating that the region was not responsive in the passive condition), the latency in the active condition is shown instead. Regions for which latency was not calculable in either the passive or active condition are not shown. Supplementary Figure S1 shows latency in both conditions for all measured brain regions (note that latency in the active condition is harder to interpret due to the smaller number of stimuli available for averaging responses and because our latency estimates require a high signal-to-noise ratio).

**Figure 3.**
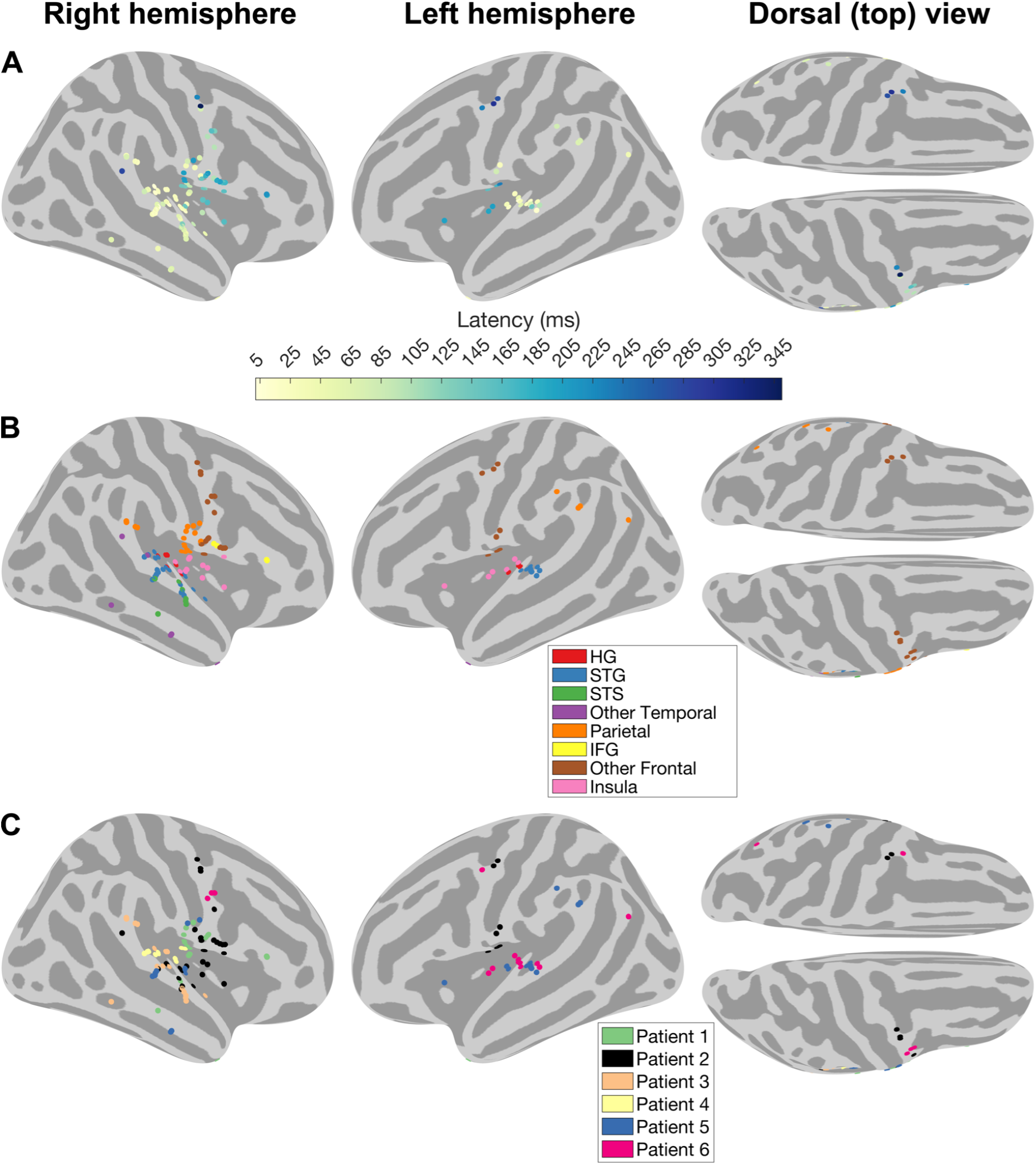
Single-channel neural response latencies in the passive condition. Panel A shows latencies for different sEEG channels across all subjects on a standard MNI brain; data are shown only for those channels for which latency was calculable in the passive condition. Panel B indicates the brain region (per Julich atlas) corresponding to each channel shown in Panel A. Different colors indicate different brain regions. Note: HG = Heschl’s gyrus, STG = superior temporal gyrus, STS = superior temporal sulcus, IFG = inferior frontal gyrus. Panel C indicates the patient ID corresponding to each channel shown in Panel A. Different colors indicate different patients.

**Figure 4.**
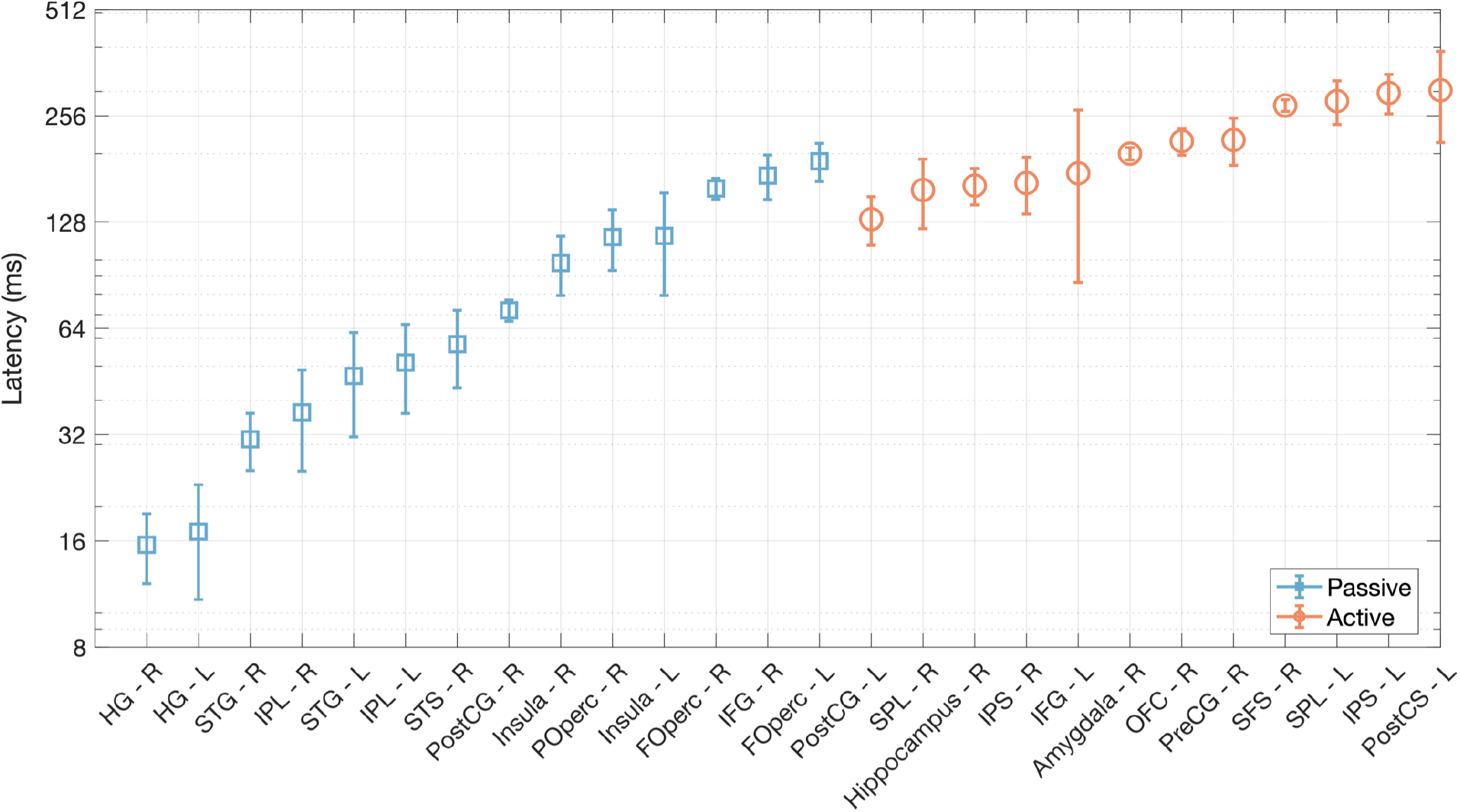
Average (over all channels across subjects) neural response latency for different brain regions in the passive (blue squares) and active (red circles) conditions. Data in the active condition are shown only for those regions that were not responsive to the stimuli in the passive condition. Error bars indicate the standard error of the mean. The different brain regions are sorted (from left to right) in the increasing order of mean latency in the passive condition; regions that were responsive only in the active condition are sorted (from left to right) in the increasing order of mean latency in the active condition. Note: Foperc = frontal operculum, HG = Heschl’s gyrus, IFG = inferior frontal gyrus, IPS = intraparietal sulcus, IPL = inferior parietal lobule, OFC = orbitofrontal cortex, POperc = parietal operculum, PostCG = Postcentral gyrus, PostCS = postcentral sulcus, PreCG = precentral gyrus, SFS = superior frontal sulcus, SPL = superior parietal lobule, STG = superior temporal gyrus, STS = superior temporal sulcus. Also note that ‘R’ indicates right hemisphere whereas ‘L’ indicates left hemisphere.

Figures 3 and 4 suggest that neural response latency is a few tens of ms in HG, several tens of ms in STG, STS, and certain parietal areas, and hundreds of ms in later parietal areas, insula, frontal cortex, hippocampus, and amygdala. Overall, there is a pattern of increasing latency in both the right [F(5, 69.07) = 11.305, p = 5.509e-08] and left [F(4, 13.895) = 4.8798, p = 0.01139] hemispheres as we ascend from auditory cortex to parietal and frontal regions.

The result that STG, STS and certain parietal areas (e.g., inferior parietal lobule; IPL) have similar latencies that are longer than the latencies in HG (Figures 3 and 4) is consistent with parallel flow of speech information dorsally and ventrally, from HG to certain parietal sites and from HG to STG and STS, respectively.

Figure 4 latency results also suggest that the following brain regions are not responsive to speech in the passive condition (latency was not calculable) but are responsive in the active condition (i.e., in the context of the speech categorization task): superior parietal lobule (SPL), hippocampus, intraparietal sulcus (IPS), amygdala, orbitofrontal cortex (OFC), precentral gyrus (PreCG), and superior frontal sulcus (SFS) in the right hemisphere, and Postcentral gyrus (PostCG), IFG, SPL, IPS, and Postcentral sulcus (PostCS) in the left hemisphere.

### Neural response amplitudes reveal regions that encode stimulus-faithful versus task-relevant representations

In addition to the latency analysis described above, we also plotted the average sEEG HGA time course for each listening condition (passive, active) to visually examine whether there may be task-related effects in the neural response amplitudes (Supplementary Figure S2). These data raise the possibility that the neural response amplitudes may differ between the active and passive listening conditions. We then directly tested for task-related effects by comparing the difference in the mean HGA amplitude between the active and passive conditions with a non-parametric null distribution to generate p-values (for analysis details, see Materials and Methods: Analysis of task-related effects and Materials and Methods: Statistical analysis). Figure 5 plots the difference between active and passive conditions in the average HGA z-score, and shows the uncorrected p-values from statistical testing for task-related effects. The brain regions that showed a task-related effect after correcting for multiple comparisons (5% FDR) are listed in Table 2.

**Table 2.** Brain regions that showed a task-related effect.

| No. | Region (hemisphere) |
| --- | --- |
| 1 | Heschl's Gyrus (Right, Left) |
| 2 | Superior Temporal Gyrus (Right, Left) |
| 3 | Inferior Parietal Lobule (Right, Left) |
| 4 | Superior Temporal Sulcus (Right) |
| 5 | Postcentral Gyrus (Right, Left) |
| 6 | Insula (Right, Left) |
| 7 | Parietal Operculum (Right) |
| 8 | Superior Parietal Lobule (Right, Left) |
| 9 | Hippocampus (Right) |
| 10 | Intraparietal Sulcus (Right, Left) |
| 11 | Inferior Frontal Gyrus (Left) |
| 12 | Amygdala (Right) |
| 13 | Orbitofrontal Cortex (Right) |
| 14 | Precentral Gyrus (Right) |
| 15 | Superior Frontal Sulcus (Right) |
| 16 | Postcentral Sulcus (Left) |

**Figure 5.**
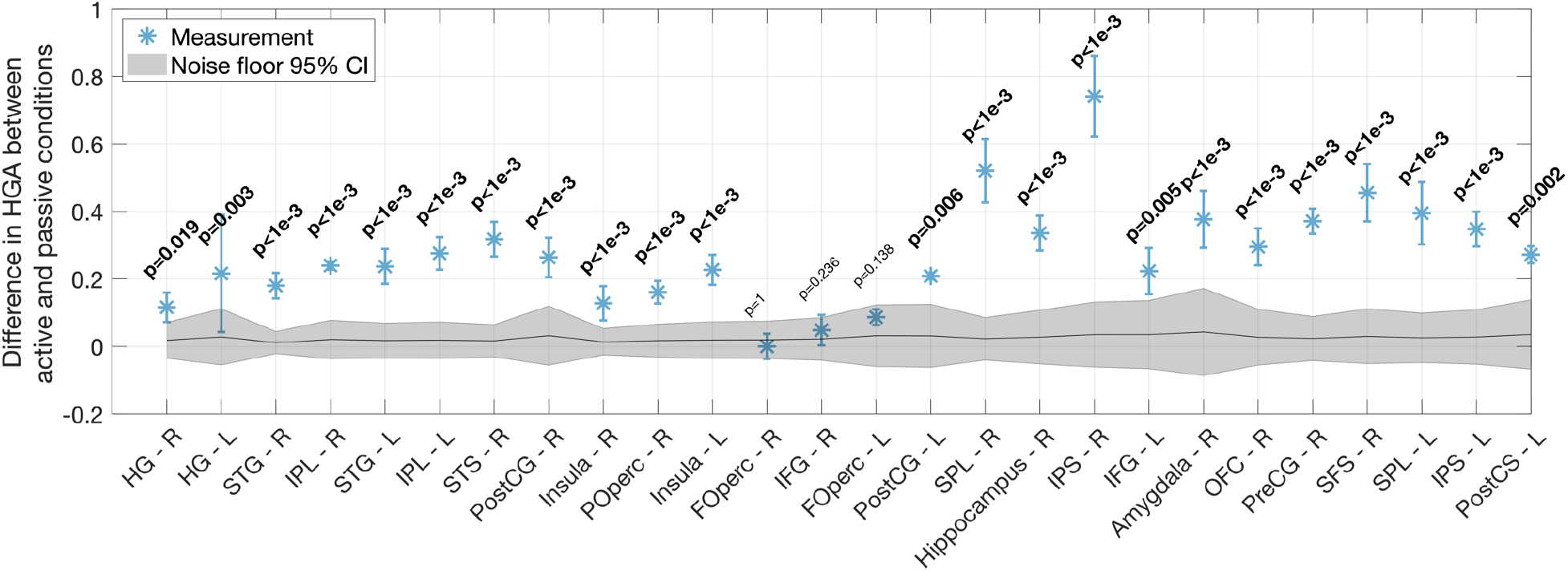
Task-related effects in different brain regions. Mean and standard error (over all channels across subjects) of the difference between active and passive conditions in the average (over time and epochs) HGA z-score relative to baseline. Also shown are uncorrected p-values for the data under the null hypothesis of zero task-related effects. Regions with statistically significant task effects at 5% FDR have bolded p-values. The different brain regions are sorted (from left to right) in the increasing order of their mean latency in the passive condition; regions that were responsive only in the active condition are sorted (from left to right) in the increasing order of mean latency in the active condition. The 95% confidence interval (CI) of the null distribution is shown as a gray patch. Note: FOperc = frontal operculum, HG = Heschl’s gyrus, IFG = inferior frontal gyrus, IPS = intraparietal sulcus, IPL = inferior parietal lobule, OFC = orbitofrontal cortex, POperc = parietal operculum, PostCG = Postcentral gyrus, PostCS = postcentral sulcus, PreCG = precentral gyrus, SFS = superior frontal sulcus, SPL = superior parietal lobule, STG = superior temporal gyrus, STS = superior temporal sulcus. Also note that ‘R’ indicates right hemisphere whereas ‘L’ indicates left hemisphere.

## Discussion

We used sEEG to quantify intracranial neural response latencies and task-related effects for spoken syllable processing in different regions of the human brain, including both sulcal and gyral structures. Our results suggest that, overall, there is a pattern of increasing latency in both the right and left hemispheres as we ascend from auditory cortex to parietal and frontal regions. Specifically, latency is a few tens of ms in HG, several tens of ms in STG, STS, and early parietal areas, and hundreds of ms in later parietal areas, insula, frontal cortex, hippocampus, and amygdala. The latency data reported in the current study for auditory cortex are of the same order of magnitude as those reported in prior intracranial (Nourski et al., 2014; Hamilton et al., 2021) and MEG/EEG (Yvert et al., 2001) studies. However, the latencies we report for forebrain areas other than auditory cortex are previously unreported to the best of our knowledge.

Our latency data are consistent with parallel streams of speech processing dorsally and ventrally, from primary auditory cortex (HG) to parietal areas and from HG to STG and STS, respectively (Figures 3 and 4). This finding is in line with dual-stream models of auditory and speech processing (Wernicke, 1874; Rauschecker, 1998; Scott, 2005; Hickok and Poeppel, 2007; Rauschecker and Scott, 2009). Some prior intracranial studies (Nourski et al., 2014; Hamilton et al., 2021) have reported parallel processing of speech within auditory cortex (specifically, that a site in STG activates in parallel with HG). Our results do not speak to this proposition as there is insufficient data to compare latencies across different anatomical sites in STG and HG. Rather, our focus is on across-region comparisons.

Overall, our latency data are consistent with hierarchical processing of speech at a macro level and parallel streams of information flow in temporal and parietal regions. Note, however, that these latency results do not imply that there is a strictly one-way flow of speech information. Indeed, there are known interactive and top-down influences along the auditory pathway from cochlea to cortex (McClelland et al., 2006; Ahveninen et al., 2006; Hickok and Poeppel, 2007; Rauschecker and Scott, 2009; Bizley and Cohen, 2013; Kemmerer, 2014).

Our combined sEEG pre-processing and latency computation procedures prevent our latency estimates from being biased by filter group delays and response smearing in time. Unlike prior approaches to estimating response latencies from intracranial data (Nourski et al., 2014; Hamilton et al., 2021), our procedures explicitly consider the impact of filter type (FIR versus IIR) and filter order (i.e., length) on the group delay of the filtered neural response and the extent of response smearing, both of which can bias latency estimates (see Supplementary Figure S3) if not properly accounted for. Taking these factors into account will ensure the best estimation of neural response latencies.

Our latency data also suggest that some brain regions are not responsive to speech during passive listening, but are responsive during active listening in the context of the speech categorization task. These regions include SPL, hippocampus, IPS, amygdala, OFC, PreCG, and SFS in the right hemisphere, and PostCG, IFG, SPL, IPS, and PostCS in the left hemisphere. To further investigate task-related effects, we compared neural response amplitudes (complementing the latency/timing analysis described above) between the passive and active conditions. We find that response amplitudes are greater in the active compared to the passive condition in several brain regions, listed in Table 2.

Our report of task effects is consistent with prior studies using fMRI that found that HG, STG, STS, IPL, PreCG, PostCG, and insula show greater activation in active versus passive listening (Jäncke et al., 1999; Hall et al., 2000; Paltoglou et al., 2009). Our results are also in line with prior intracranial (Mesgarani and Chang, 2012; O’Sullivan et al., 2019), MEG (Woldorff et al., 1993; Kauramäki et al., 2012; Ding and Simon, 2012), and EEG (Choi et al., 2014; Viswanathan et al., 2019) studies that suggest that attention modulates responses even in early auditory cortical areas (HG and STG). Furthermore, our findings are also consistent with increased engagement of sensorimotor areas like left-hemisphere IFG during phoneme categorization tasks (Hickok et al., 2011; Skipper et al., 2017). Indeed, in general, the task effects we report in the present study can be attributed to a combination of several perceptual processes, including attention, reading, and decision masking in speech categorization. However, they are likely not attributable to low-level processing of differences in the visual stimuli between conditions or to button-press-related motor planning signals, as described below.

A caveat in our report of task effects is that the visual stimuli are different in the active and passive conditions. However, we do not report results from visual cortex and it is unlikely that the aggregate task-related effects we see in other brain regions reflect low-level processing of differences in the visual stimuli. Some of the task effects we report could be due to higher level processing (e.g., reading) of the visual stimuli associated with the task. However, all of our analyses are time-locked to the auditory stimuli. Moreover, the effects we report are sustained over time and not transient. Thus, any differences we observe in the neural response amplitudes between active and passive conditions likely reflect a true task effect related to syllable categorization. Furthermore, the task effects we find are also unlikely to reflect button-press-related motor planning or motor action signals. This is because the median time at which the button presses occurred relative to the visual cue asking subjects to choose between /b/ and /p/ in the active condition is 1.52 seconds (the median absolute deviation is 0.8396 seconds), which is longer than the duration of our analysis epochs (450 ms post stimulus onset).

A more general caveat of our study is that because sEEG affords sparse (versus whole-brain) electrode coverage, the absence of a region in our latency and task-effects reports does not necessarily imply that that region does not participate in speech processing. Rather, this could simply be due to insufficient sEEG coverage in that region. For example, in the present study, coverage is sparser in the left (versus right) hemisphere (Figure 3). Consequently we do not report latency and task-related effects in some left-hemisphere regions like STS. This should be explored in future intracranial studies.

## Acknowledgements

This study was supported by National Institutes of Health grants F32DC020649 (to VV), T32DC11499 (institutional support to VV), R21DC019217 (to TJA and LLH), and R01DC013315 (to TJA).

