## Supplementary Figures for "Intracranial Mapping of Response Latencies and Task Effects for Spoken Syllable Processing in the Human Brain"

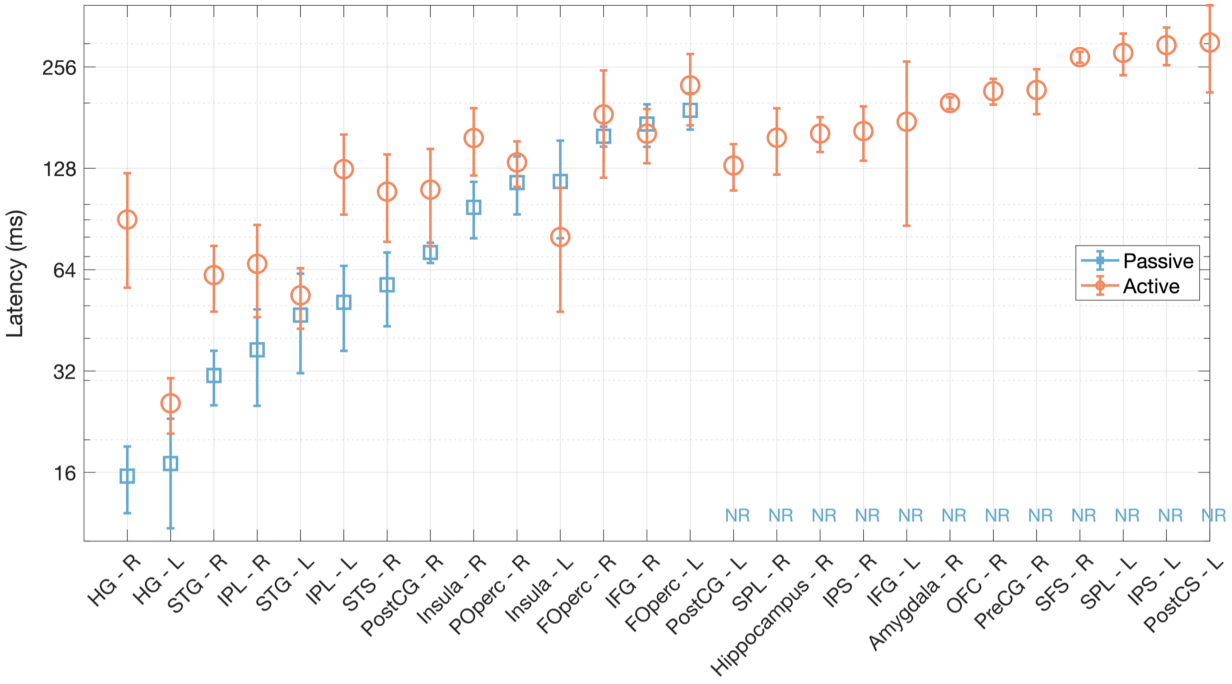


**Supplementary Figure S1.** Average (over all channels across subjects) neural response latency for different brain regions in the passive (blue squares) and active (red circles) conditions. Error bars indicate the standard error of the mean. The different brain regions are sorted (from left to right) in the increasing order of their mean latency in the passive condition; regions that were responsive only in the active condition are sorted (from left to right) in the increasing order of mean latency in the active condition. Regions for which latency was not calculable (i.e., the region was not responsive to the speech stimuli) in the passive condition are indicated as NR (Not Responsive). Note: FOperc = frontal operculum, HG = Heschl's gyrus, IFG = inferior frontal gyrus, IPS = intraparietal sulcus, IPL = inferior parietal lobule, OFC

= orbitofrontal cortex, POperc = parietal operculum, PostCG = Postcentral gyrus, PostCS = postcentral sulcus, PreCG = precentral gyrus, SFS = superior frontal sulcus, SPL = superior parietal lobule, STG = superior temporal gyrus, STS = superior temporal sulcus. Also note that ‘R’ indicates right hemisphere whereas ‘L’ indicates left hemisphere.

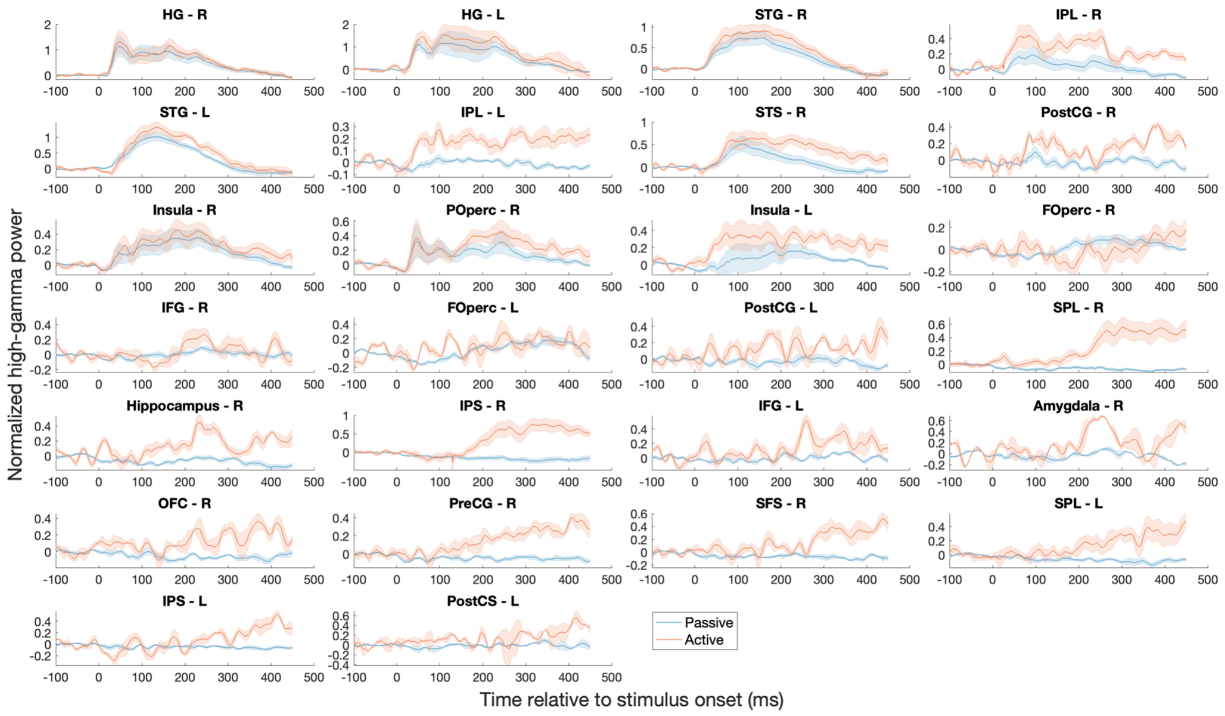


**Supplementary Figure S2.** Neural response time courses in different brain regions. Mean time courses (HGA z-score relative to baseline, averaged over all epochs, responsive channels, and subjects) are shown separately for the active (red curve) and passive (blue curve) conditions. Shaded error bars indicate the standard error across channels and subjects. The different brain regions are sorted (from left to right, and top to bottom) in the increasing order of their mean latency in the passive condition; regions that were responsive only in the active condition are sorted (from left to right, and top to bottom) in the increasing order of mean latency in the active condition. Data are shown only for those brain regions for which latency was calculable (i.e., the region was responsive to the speech stimuli) in either the passive or active condition. Note: FOperc = frontal operculum, HG = Heschl’s gyrus, IFG = inferior frontal gyrus, IPS = intraparietal sulcus, IPL = inferior parietal lobule, OFC = orbitofrontal cortex, POperc = parietal operculum, PostCG = Postcentral gyrus, PostCS = postcentral sulcus, PreCG = precentral gyrus, SFS = superior frontal sulcus, SPL = superior parietal lobule, STG = superior temporal gyrus, STS = superior temporal sulcus. Also note that ‘R’ indicates right hemisphere whereas ‘L’ indicates left hemisphere.

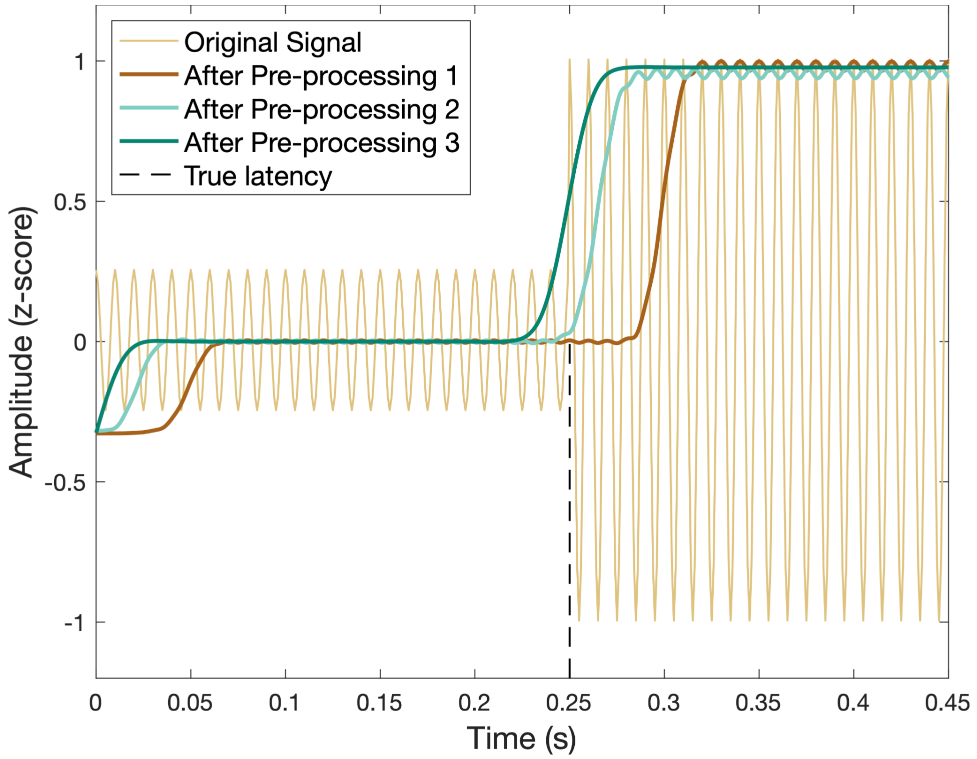


**Supplementary Figure S3.** Simulated effect of different pre-processing approaches on neural response group delay and latency estimates. A 100 Hz sinusoid ("original signal"; shown in light brown) with a baseline period (0 to 0.25 s) of relatively low amplitude and subsequent response period (0.25 s onwards) was subjected to three different pre-processing strategies. In "Pre-processing 1", the signal was filtered in the high-gamma band (75-150 Hz) using a linear-phase causal FIR filter of order 68 ms (group delay = 34 ms), then half-wave rectified and low-pass filtered between 0 and 30 Hz using a linear-phase non-negative causal FIR filter constructed using a Hanning window of length 33 ms (group delay = 16 ms) to extract the high-gamma-band envelope (i.e., high-gamma activity or HGA; shown in dark brown). In "Pre-processing 2", the original signal was filtered in the high-gamma band using a zero-group-delay (non-causal) FIR filter (order = 68 ms), then half-wave rectified and low-pass filtered between 0 and 30 Hz using the same causal FIR filter as in Pre-processing 1 to extract HGA (shown in light green). "Pre-processing 3" used the same steps as the other two approaches, but with zero-group-delay (non-causal) filtering throughout (result shown in dark green). In the case of Pre-processing 2 (the approach used in the main analyses of the present study), the time point of initial deviation of the pre-processed signal is 0.25 s, which is equal to the true latency of the original signal (i.e., the time at which the signal amplitude deviates from baseline activity). In the case of the other two pre-processing strategies, the accumulated group delays and signal smearing in time from filtering can cause latency to be incorrectly estimated.
